# Tartrazine clears live cells while preserving viability at high refractive indices and osmolality

**DOI:** 10.64898/2026.04.09.717314

**Authors:** Xuandi Hou, Sa Cai, Han Cui, Zhongyu Liu, Su Zhao, Ling-Yi Zhang, Ani Baghdasaryan, Victoria Crunkleton, Mark L. Brongersma, Guosong Hong

## Abstract

Tissue-clearing techniques have transformed optical imaging of fixed specimens, yet their application to living systems remains limited by toxicity and removal of key tissue components. We recently demonstrated that absorbing molecules such as tartrazine can reversibly render live mouse skin transparent. Subsequently, it was reported that isotonic protein solutions can achieve *ex vivo* and *in vivo* cellular clearing. However, discrepancies remain regarding the optimal refractive index (RI) for live-cell clearing and the impact of elevated osmolality on cell viability. Here, using cultured mammalian cells, we systematically examine the dependence of optical contrast on medium RI and the effects of hyperosmolality. We find that, contrary to the recent report of an optimal RI of 1.36∼1.37 for suspended cells, densely-packed adherent cells exhibit a monotonic decrease in phase contrast up to an RI of 1.41 with tartrazine. Moreover, even under highly hyperosmotic conditions (∼1200 mOsm/kg), cultured cells exhibit minimal deformation and negligible loss of viability for up to 30 min in the clearing solution. These results demonstrate that tartrazine enables effective live-cell clearing at RI up to 1.41 while preserving viability under elevated osmolality, and motivate future studies to define optimal conditions for *in vivo* optical clearing.

**Graphical Abstract:** 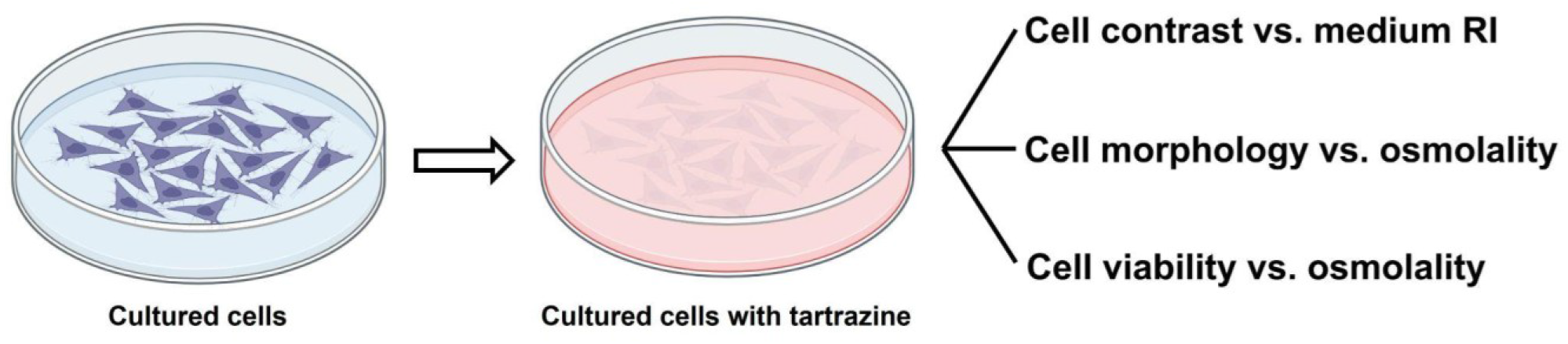

## Introduction

Light is essential to biology and medicine, enabling researchers to explore complex biological processes and to diagnose and treat a wide range of medical conditions with precision and efficacy.^1,2^ However, optical imaging and light delivery in deep tissues remain challenging due to limited light penetration, which hinders volumetric interrogation of biological systems.^3,4^ This limitation is primarily caused by photon scattering and absorption, with scattering playing the dominant role.^5^ Therefore, developing strategies to reduce scattering is critical for advancing deep-tissue imaging and improving light-based biological interventions.

Light scattering arises from differences in refractive indices (RI) between low-RI aqueous components (e.g., the interstitial fluid and cytosol) and high-RI lipid/protein components (e.g., the plasma membrane, myelin, and extracellular matrix).^6^ A prevailing view holds that water-rich cytoplasm and interstitial fluid have RI of 1.35∼1.37, in contrast to lipid-rich plasma membranes and intracellular organelles, which exhibit higher RIs of 1.41∼1.46.^6–10^ Based on this understanding of RI heterogeneity within tissues, conventional tissue-clearing technologies minimize light scattering and thus render tissues transparent by removing water or lipids, resulting in a more optically homogeneous residual medium.^9–11^ However, these methods are incompatible with live tissues due to toxicity and removal of water or lipids, both of which are essential for sustaining life.^9,11^ To address these challenges, our lab and many other labs recently demonstrated that strongly absorbing dye molecules, such as tartrazine, ampyrone, fluorescein, and indocyanine green, can homogenize the RI between aqueous and non-aqueous components, thereby achieving optical transparency in the tissues of live animals.^12–17^ Since our original report,^12^ this dye-based *in vivo* tissue clearing approach has been rapidly adopted by laboratories worldwide, enabling a range of new applications.^18–38^

More recently, a study reported optical clearing of live cells both *ex vivo* and *in vivo* using high concentrations of bovine serum albumin (BSA).^39^ This work suggested that the optimal RI for live-cell clearing lies within a narrow range of 1.36∼1.37, in contrast to the prevailing view that effective hydrophilic clearing requires raising the RI of the medium toward that of lipid-rich cellular components (1.41∼1.46).^6–10^ Notably, this lower optimal RI was primarily inferred from optical transmission measurements in suspended cells lacking sufficient extracellular matrix (ECM).^39^ Furthermore, because BSA is a large macromolecule (molecular weight ∼66.5 kDa) that does not readily cross the plasma membrane, it remains osmotically active in the extracellular space. Based on this property, the study posited that clearing agents must be isotonic to maintain compatibility with live cells during imaging. Here, we provide evidence that challenges these conclusions. Using tartrazine as a model small-molecule clearing agent, we show that optical contrast in densely-packed, adherent human embryonic kidney (HEK) cells decreases monotonically with increasing medium RI up to 1.41, without evidence of an optimal range at 1.36∼1.37. In addition, despite the high osmolality associated with tartrazine solutions, cells exhibit minimal shrinkage under hypertonic conditions and remain viable over at least 30 min.

We attribute these discrepancies primarily to differences between suspended individual cells and densely-packed adherent cellular systems, with the latter more closely recapitulating *in vivo* conditions due to the presence of ECM and cell-cell interactions. Moreover, we argue that the requirement for isotonicity is not universal; instead, it depends on the membrane permeability of the clearing agent and the tolerance of the specific cell type and its surrounding milieu. Specifically, our results show that hyperosmotic tartrazine is tolerated by adherent HEK293 cells for at least 30 min, likely due to intrinsic regulatory mechanisms that preserve viability. These findings highlight the need for future studies to resolve these discrepancies and to guide the rational design of more effective and biocompatible *in vivo* clearing agents.

## Results

### Optical characterization of tartrazine and gelatin solutions

We set out to demonstrate that a mixture of tartrazine and gelatin, at effective concentrations, can reduce the scattering of silica nanospheres (RI = 1.43) suspended in water and thereby achieve optical transparency. Specifically, an aqueous suspension of 1-μm diameter silica nanospheres (10 mg/mL) in water exhibits pronounced opacity, which is not alleviated by the addition of 5.8% w/w gelatin alone. Similar to other biocompatible and water-soluble polymers (e.g., agarose and polyvinyl alcohol) used in our previous studies,^12,13^ this low concentration of gelatin serves to stabilize tartrazine at its effective concentration while minimally perturbing the intrinsic optical properties of the medium (**Fig. 1A**). In contrast, when tartrazine is added to this mixture at a concentration of 0.47 M, it effectively reduces scattering, resulting in significant optical transparency in the red spectrum (**Fig. 1A**). The ability of the tartrazine/gelatin mixture to enhance transparency in this scattering system is further confirmed by transmission spectra (**Fig. 1B**). We attribute this transparency effect to tartrazine absorption near 420 nm, which modulates the background RI to ∼1.41 at longer wavelengths, thereby more closely matching the RI of the silica particles (**Fig. 1C&D**). Despite its strong absorption near 420 nm, tartrazine exhibits minimal absorption beyond 600 nm (**Fig. 1E**), where RI matching can reduce scattering without introducing additional absorption. Due to gelatin’s minimal absorption in the visible spectrum (**Fig. 1E**), its presence in the tartrazine solution only causes minor changes to the primary absorption peak of tartrazine at ∼420 nm (**Fig. 1F**), as well as to the real and imaginary components of the RI measured by spectroscopic ellipsometry (**Fig. 1C&D**).

**Figure 1.**
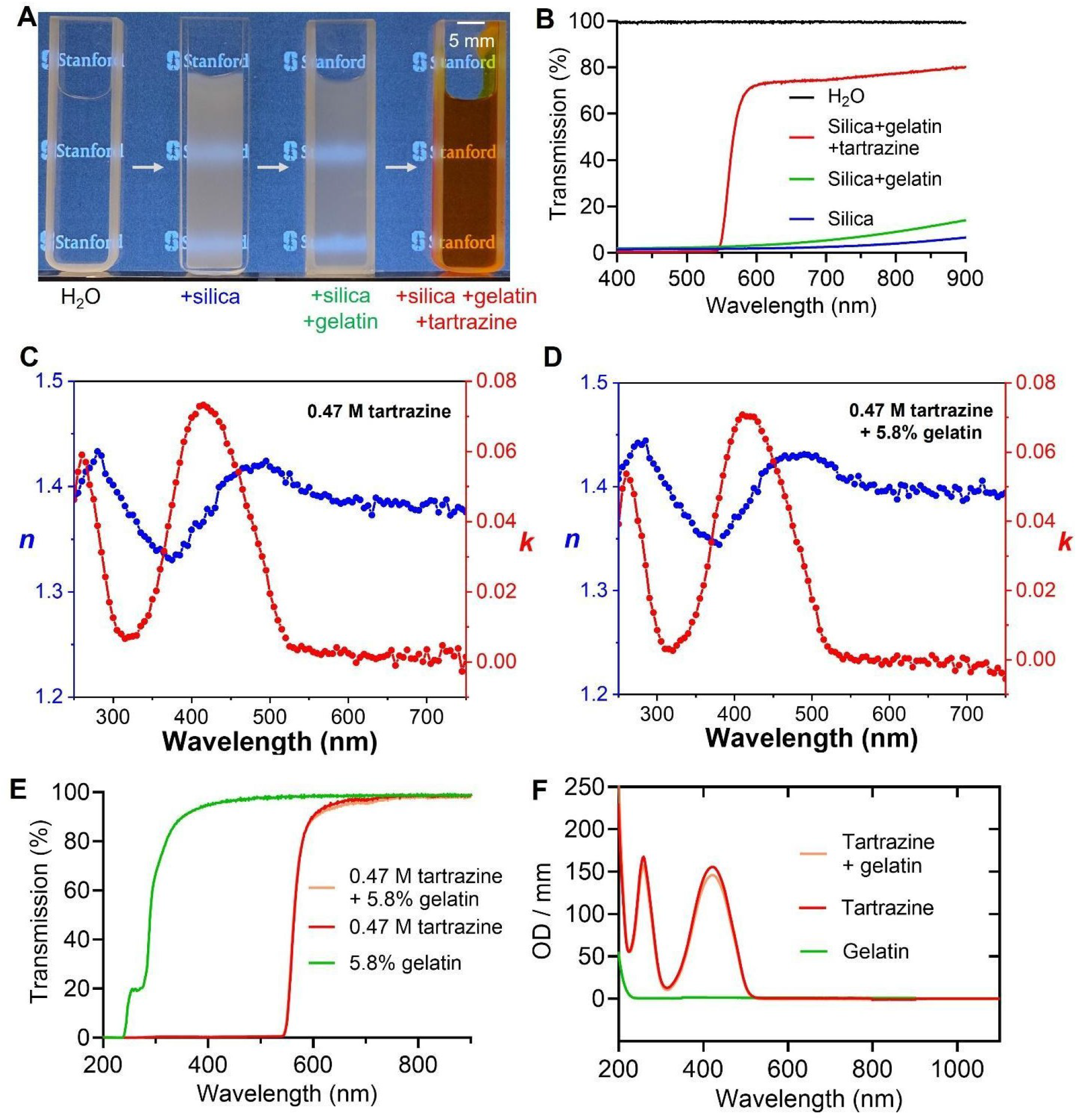
Optical characterizations of tartrazine and gelatin solutions. (**A**) Brightfield transmission images of different media in cuvettes with a 1-mm path length. From left to right: pure water, 1-μm colloidal silica particles (10 mg/mL) suspended in water, the same silica particles suspended in an aqueous solution containing 5.8% w/w gelatin, and the same silica particles suspended in an aqueous solution containing 5.8% w/w gelatin and 0.47 M tartrazine. (**B**) Transmission spectra of the samples in **A**. (**C&D**) Real (*n*) and imaginary (*k*) components of the refractive index of the 0.47 M tartrazine solution (**C**) and 0.47 M tartrazine + 5.8% w/w gelatin solution (**D**), showing minimal changes in the ellipsometry data of tartrazine due to the presence of gelatin. (**E**) Transmission spectra of tartrazine, gelatin, and tartrazine + gelatin solutions, all without silica particles and measured over a 1-mm path length. (**F**) Absorption spectra of gelatin, tartrazine, and tartrazine + gelatin solutions, showing minimal contribution of gelatin to the absorption peaks of tartrazine. These solutions were obtained by diluting the corresponding solutions in **E** by a factor of 9.4 to avoid OD saturation on the UV-Vis spectrometer.

### Tartrazine/gelatin solution clears live cells with increasing transparency up to an RI of 1.41

We next sought to demonstrate the effect of the tartrazine/gelatin solution on the optical contrast of live cell cultures under microscopic imaging. RI differences among various intracellular and extracellular components, such as the plasma membrane, organelles, cytoplasm, interstitial fluid, and ECM, are the primary sources of contrast in differential interference contrast (DIC) microscopy of cultured HEK cells (**Fig. 2A**). It is generally accepted that water-rich cytoplasm and interstitial fluid exhibit lower RIs of 1.35∼1.37, whereas lipid-rich plasma membranes, organelles, and ECM protein fibers possess higher RIs of 1.41∼1.46.^6–10^ Upon exposure to increasing tartrazine concentrations (and thus increasing RI), densely packed HEK cells cultured in Petri dishes exhibited a progressive loss of optical contrast within 2 min of solution application. At 0.47 M tartrazine (RI = 1.41), the loss of contrast was most pronounced, rendering cellular boundaries and intracellular features nearly imperceptible under microscopy due to RI matching (**Fig. 2B**; **Supplementary Movie 1**). Quantitative line profile analysis of representative cell images confirmed reduced intensity fluctuations (**Fig. 2C**), corresponding to decreased cellular contrast (**Fig. 2D**). Importantly, the optical contrast of densely-packed, adherent HEK cells decreased monotonically with increasing medium RI (**Fig. 2D & Fig. S1A**). As a control experiment, the addition of gelatin up to 5.8% w/w only slightly increases the RI of the solution (**Fig. S1B**). Specifically, aqueous solutions with increasing tartrazine concentrations exhibit a progressively deeper red color, while remaining highly transparent in the red channel (**Fig. S2**). Taken together, these results indicate that increasing tartrazine concentrations, and the associated monotonic rise in RI up to 1.41, progressively diminish cellular contrast in DIC microscopy, while tartrazine’s transmission above 600 nm still permits microscopic cellular imaging.

**Figure 2.**
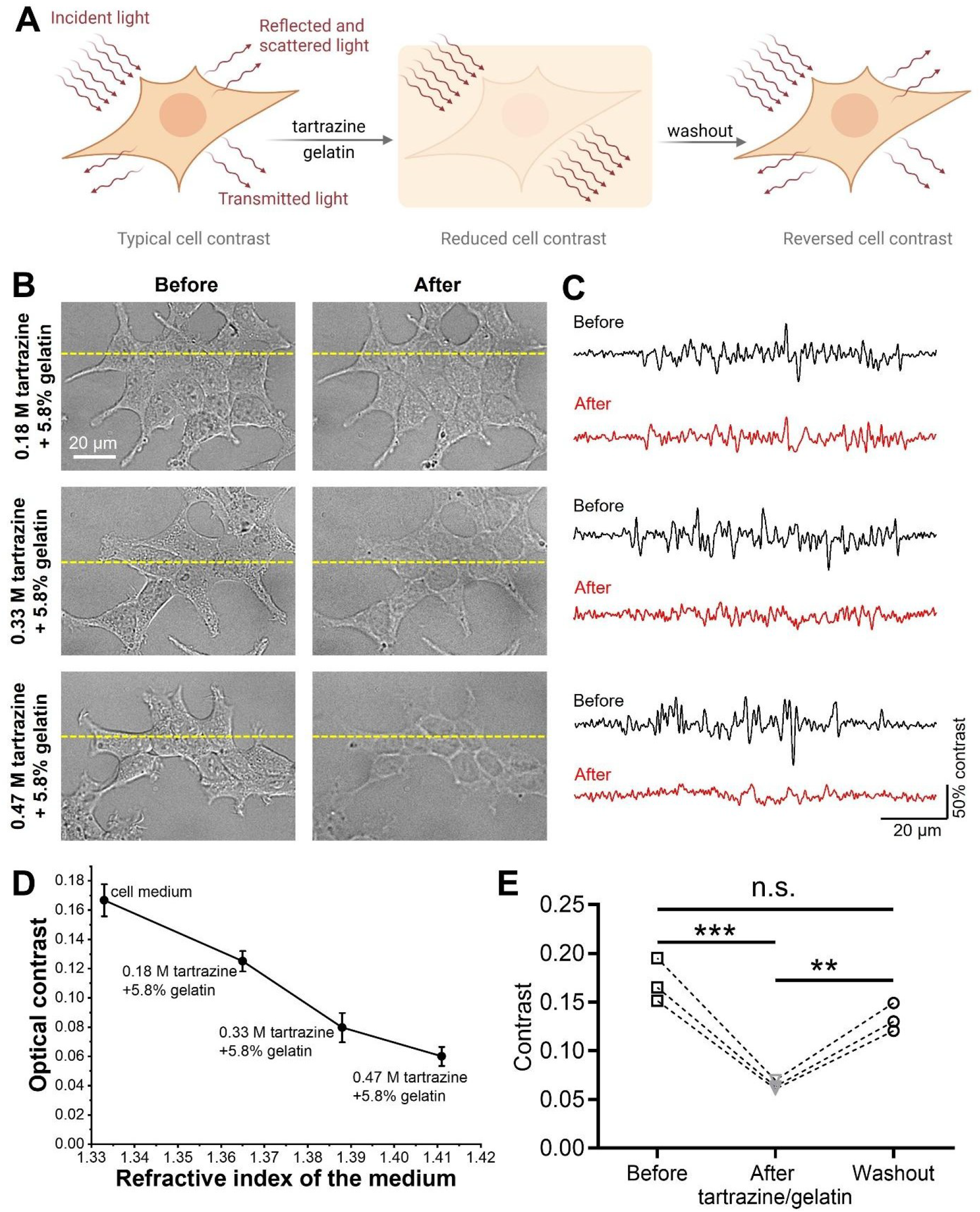
Cultured cells become progressively more transparent as RI increases up to 1.41. (**A**) Schematic illustration depicting the procedure of reducing and reversing optical contrast of cultured cells. (**B**) DIC images of HEK cells before (left) and after (right) exposure to solutions containing different concentrations of tartrazine and 5.8% w/w gelatin. (**C**) Intensity profiles along the yellow dashed lines in **B** before and after adding the tartrazine/gelatin solution. The grayscale intensity is mapped to a 0–100% range based on the minimum and maximum of each image. (**D**) Quantitative optical contrast of HEK cells as a function of the medium RI. Data in **D** is presented as mean ± SD for n = 6 biological replicates. (**E**) Quantitative optical contrast of HEK cells before exposure to the tartrazine/gelatin solution, during exposure, and after washing out the tartrazine/gelatin solution with tartrazine-free medium. Experiments in **E** were performed in n = 3 biological replicates. n.s., non-significant (p ≥ 0.05), **p < 0.01, ***p < 0.001, ****p < 0.0001, one-way ANOVA.

The observed loss in optical contrast in cultured HEK cells is reversible. Specifically, replacing the tartrazine-containing medium with that free of tartrazine and gelatin largely restores the original cellular contrast (**Fig. 2E**). This reversible phenomenon is consistent with similar observations in live mice.^12,13,32,38^ In summary, these findings reveal that the reduced optical contrast of cultured cells aligns with, and underlies, the macroscopic tissue transparency observed in live animals.^12,13^ Notably, we observed a monotonic increase in the transparency of densely-packed adherent cells with increasing medium RI, extending up to an RI of 1.41, in contrast to prior reports identifying an optimal RI of 1.36∼1.37 in suspended cells.^39^ These results indicate that effective reduction of optical contrast in cultured cells is achieved by tuning the extracellular medium RI toward that of lipid-rich cellular components (1.41∼1.46), rather than matching the RI of the cytoplasm (1.35∼1.37).

### Tartrazine/gelatin solution induces minimal cell shrinkage despite high osmolality

We next asked whether tartrazine, at concentrations effective for optical clearing, alters the size and morphology of these “cleared” cells. This question arises from the measured osmolality of 1220 mOsm/kg for tartrazine at its effective concentration (0.47 M). Due to the hyperosmotic condition, exposure of HEK cells to 0.47 M tartrazine induces a ∼20% reduction in cell area within 2 min, accompanied by a concurrent loss of plasma membrane contrast, as expected from the optical clearing effect (**Fig. 3A-C**; **Supplementary Movie 2**). This ∼20% reduction in cell area is consistent with literature reports showing that cultured cells with comparable, weak adhesion strength undergo a similar (15∼25%) decrease in size upon ∼2 min exposure to hypertonic media.^40–42^

**Figure 3.**
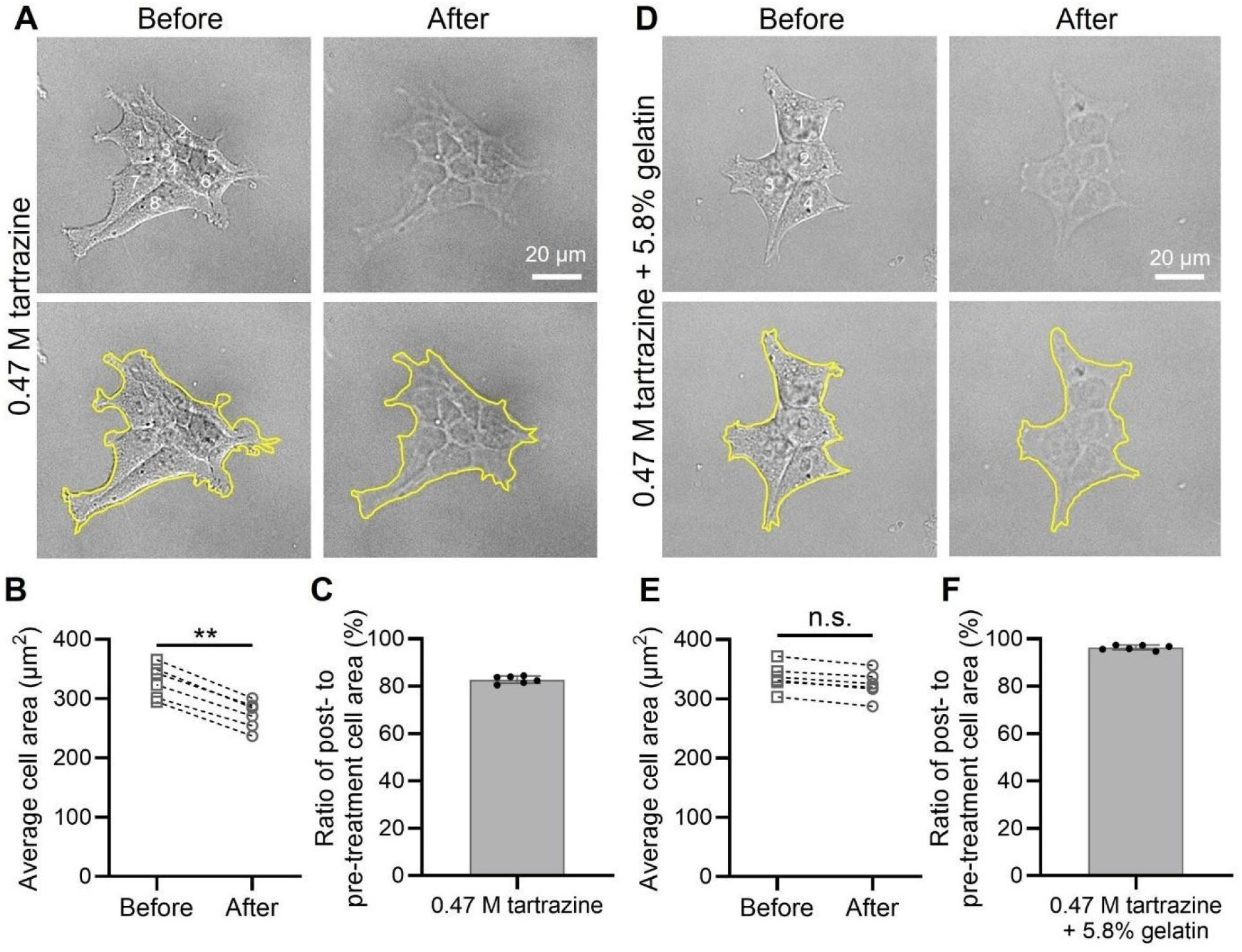
Cultured cells undergo minimal shrinkage while achieving transparency under high osmolality. (**A**) HEK cells immersed in a solution containing 0.47 M tartrazine alone exhibit ∼20% cell shrinkage. (**B**) Quantification of the average cell area following tartrazine treatment in **A** for n = 6 biological replicates. (**C**) Quantification of the percent cell area change following tartrazine treatment in **A**. (**D**) HEK cells immersed in a solution containing 0.47 M tartrazine and 5.8% w/w gelatin exhibit negligible cell shrinkage. (**E**) Quantification of the average cell area following tartrazine/gelatin treatment in **D** for n = 6 biological replicates. (**F**) Quantification of the percent cell area change following tartrazine/gelatin treatment in **D**. n.s. indicates non-significant differences (p ≥ 0.05), **p < 0.01, as determined by unpaired two-tailed t-tests. Data in panels **C** and **F** are presented as mean ± SD for n = 6 biological replicates.

More surprisingly, the addition of increasing concentrations of gelatin progressively attenuates cellular shrinkage, while preserving the clearing effect (**Fig. S3&S4**; **Fig. 3D-F**; **Supplementary Movie 1**). Specifically, at a gelatin concentration of 5.8% w/w, cultured cells exhibit reduced membrane contrast without a significant decrease in cell size (**Fig. 3D-F**). Gelatin was selected based on our prior work using water-soluble polymers (e.g., agarose and polyvinyl alcohol) as chemical stabilizers for tartrazine solutions for *in vivo* clearing.^12,13^ A salient advantage of gelatin is that it maintains the medium in a liquid state and offers well-documented biocompatibility for cell culture.^43^ Notably, gelatin provided benefits beyond stabilization, significantly mitigating hyperosmotic tartrazine-induced cellular shrinkage (**Fig. 3D-F**; **Fig. S3&S4**) with minimal impact on the optical properties of the tartrazine solution (**Fig. 1C-F**; **Fig. S1&2**) or its clearing effect (**Fig. 2&3**). We attribute the ability of gelatin to fully prevent cellular shrinkage to its increased viscosity and inherent tissue-supportive properties,^44–46^ rather than to any reduction in the nominal osmolality of the tartrazine solution, as the 0.47 M tartrazine/5.8% gelatin mixture still exhibits a high measured osmolality of 1288 mOsm/kg. Taken together, these experiments demonstrate that hyperosmolality, even at levels of ∼1200 mOsm/kg, does not necessarily result in shrinkage of live cells.

### Tartrazine/gelatin solution preserves cell viability despite high osmolality

Lastly, we sought to determine the effect of hyperosmotic tartrazine exposure on cell viability. Specifically, we conducted *in vitro* evaluations of cell viability following hyperosmotic tartrazine treatment (∼1200 mOsm/kg) at clearing concentrations. These assessments included live/dead cell staining, the MTT assay, and flow cytometry following incubation with and after washout of the clearing agents, aimed at evaluating their delayed impact on cellular health.

We chose a 30-min evaluation window because *in vivo* transient transparency in live mice is typically maintained over this duration, providing sufficient time for dynamic imaging before the tissue is returned to its native opaque state.^12–14^ Specifically, fluorescent live/dead staining revealed minimal cell death after 30 min of continuous cell incubation with 0.47 M tartrazine and 5.8% gelatin. Furthermore, even exposure to 0.47 M tartrazine alone (i.e., without gelatin) for 30 min resulted in viable cells comprising ∼90% of the total cell population (**Fig. 4A&B**). Notably, in both cases the cells gradually become less spread and more rounded over 30 min. We attribute this change to a gradual loss of cell-substrate adhesion rather than osmolality-induced shrinkage, as the latter would be expected to occur on a much shorter timescale than observed here (**Fig. 3**).^40,41^ The high viability confirmed by live/dead assays indicates that this morphological change results in minimal cell damage after 30 min of continuous incubation in tartrazine-containing medium. Such rounding is commonly observed in weakly adherent cells, such as HEK293 cells studied here, where loss of adhesion does not necessarily indicate cell death.^47^ In contrast, the positive control for cytotoxicity (20% DMSO) reduced the fraction of viable cells to <5% within 10 min. We also observed a small but statistically significant decrease in cell viability in the 30-min iodixanol-treated group (96%), which was identified as an alternative clearing agent for *in vitro* cell studies,^48^ compared to the group treated with 0.47 M tartrazine and 5.8% gelatin (98%) (**Fig. S5**). Together, these results support the overall biocompatibility of tartrazine as an *in vivo* tissue-clearing agent, consistent with its status as an FDA-approved food dye with a well-established safety profile, including a high median lethal dose (LD50) exceeding 12 g/kg body weight in mice.^49,50^ In addition, these findings highlight the utility of incorporating biocompatible polymers as additives to enhance both the biosafety and chemical stability of tartrazine, in agreement with our previous *in vivo* observations.^12,13^

**Figure 4.**
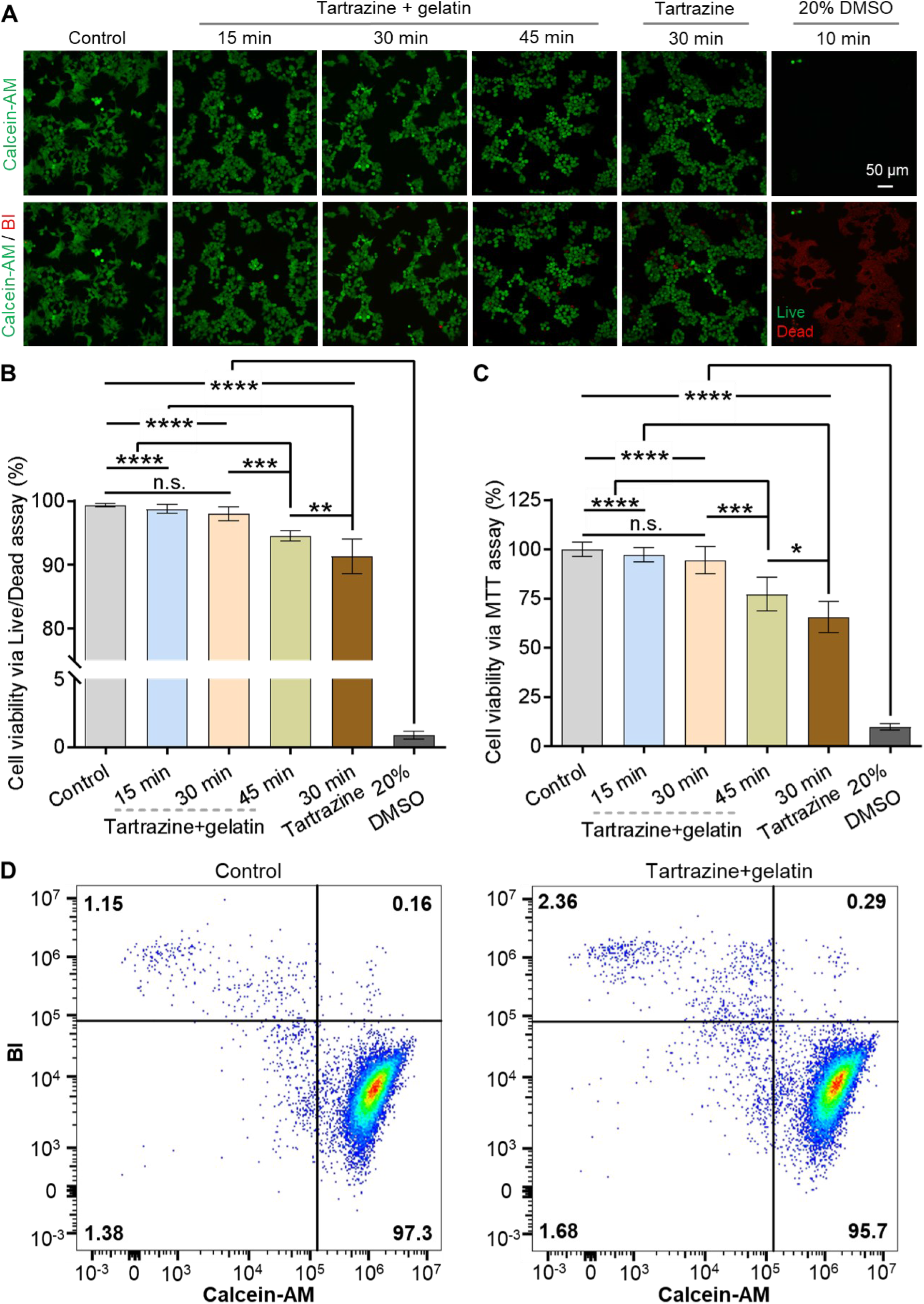
Cultured cells remain viable in tartrazine clearing solutions for at least 30 min. (**A**) Representative live/dead staining images of HEK cells incubated with 0.47 M tartrazine + 5.8% gelatin for 15, 30, and 45 min, 0.47 M tartrazine alone for 30 min, or 20% DMSO (positive control for cell toxicity) for 10 min. Green and red colors indicate live and dead cells, respectively. (**B**) Quantified cell viability of each group after incubation based on live/dead staining results. (**C**) Quantified cell viability of each group after incubation based on the MTT assay. n.s., non-significant (p ≥ 0.05), *p < 0.05, **p < 0.01, ***p < 0.001, ****p < 0.0001, one-way ANOVA. Data represent the mean ± SD for **B** and **C** without excluding any outliers. For each condition in **B&C**, *n* = 6 biological replicates. (**D**) Analysis of live and dead cell subpopulations in the control condition (without treatment) and the experimental condition containing 0.47 M tartrazine + 5.8% gelatin by flow cytometry. Numbers in each quadrant indicate the percentage of gated cells corresponding to each viability state.

Besides live/dead staining, we also performed the MTT assay to evaluate cell metabolic activity. The MTT results showed minimal changes in metabolic activity for cells treated with 0.47 M tartrazine and 5.8% gelatin compared to the tartrazine-free control. By contrast, cells treated with 0.47 M tartrazine alone exhibited a significant reduction in metabolic activity to ∼70% of control levels (**Fig. 4C**). Furthermore, we evaluated delayed biocompatibility following 30-min tartrazine/gelatin treatment to assess its suitability for potential longitudinal imaging studies. To this end, we performed flow cytometry on cells harvested from continuous culture 48 h after a 30-min exposure to 0.47 M tartrazine supplemented with 5.8% gelatin. Cells were labeled with calcein-AM and BOBO-3 iodide (BI) prior to analysis. Remarkably, cells treated with 0.47 M tartrazine and 5.8% gelatin for 30 min demonstrated viability comparable to that of control, untreated cells, indicating sustained biocompatibility with negligible delayed bioeffects (**Fig. 4D**).

We next sought to extend the duration of tartrazine/gelatin treatment to determine the time point at which significant cellular toxicity emerges. We found that extending the exposure time to 45 min, followed by washout and contrast reversal, led to a statistically significant decrease in viable cells even in the presence of 5.8% w/w gelatin (**Fig. 4B&C**). However, gelatin still provided a protective effect: even at the 45-min time point, cell viability remained substantially higher (∼95%) than that of cells exposed to tartrazine alone for only 30 min (∼90%). Taken together, these findings, which are based on multiple complementary viability assays, further confirm the overall safety of tartrazine as well as the beneficial role of gelatin despite the high osmolality of ∼1200 mOsm/kg for these tartrazine-based clearing solutions.

## Discussion

This work establishes three major findings. First, using tartrazine as a representative clearing agent, we demonstrate that live-cell transparency increases monotonically with an increasing medium RI up to 1.41. Second, despite the high osmolality of tartrazine-based clearing solutions at effective concentrations (∼1200 mOsm/kg), these conditions induce only minimal deformation in cultured cells and, in the presence of the stabilizing polymer, gelatin, can clear cells without any measurable shrinkage. Third, under these same hypertonic conditions, cultured cells maintain high viability (>90%) for exposure durations of up to 45 min, providing a sufficient time window for imaging applications.

These findings contrast with a recent report of live-cell clearing using BSA, in which an optimal RI of 1.36∼1.37 was identified based on measurements in suspended cells.^39^ In addition, that study emphasized the necessity of maintaining physiological osmolality (∼300 mOsm/kg), a requirement that is well justified for highly sensitive cell types such as neurons, but may be overly restrictive for other cell types, including those examined here.

We attribute these discrepancies to differences in extracellular components and their associated optical properties between suspended cells (in the recent report^39^) and densely-packed adherent cells (in this paper). In particular, collagen, a principal component of the ECM, has a reported RI of ∼1.47 and represents a major source of light scattering in adherent cells and three-dimensional tissues.^6–10^ Accordingly, a clearing medium with an RI of ∼1.41 provides improved RI matching to collagen-based scatterers while remaining reasonably close to the average cytoplasmic RI (1.36∼1.37). Furthermore, the ECM is known to buffer osmotic stress through its mechanical support and cell-matrix interactions,^51^ offering a plausible explanation for the minimal shrinkage observed in adherent cells under hypertonic clearing conditions. Finally, the high viability of cultured cells (∼95% after 45 min exposure) despite significantly elevated osmolality can be attributed to the combined effects of ECM-enabled osmotic buffering,^51^ the established biocompatibility of tartrazine,^49,50,52^ and the additional stabilizing role of gelatin.^44–46^

This work also provides several broader insights. First, the role of osmolality in determining the *in vivo* safety of clearing agents should be evaluated in a tissue-specific context. For example, in skin, multiple studies have demonstrated the safe use of hyperosmotic agents such as polyethylene glycol and glycerol,^53–57^ attributable in part to the osmotic buffering capacity of the collagen-rich ECM surrounding live skin cells.^51^ The tolerance of skin and mucosal tissues to hyperosmolar environments is further reflected in the high osmolality of certain topical products (often >6000 mOsm/kg)^58^ and dietary exposures.^59^ Notably, the World Health Organization recommends an upper safety limit of ∼1200 mOsm/kg for substances in direct contact with live mucosal tissues.^58^ Our findings are consistent with this threshold, supporting a broader permissible range of osmolality for *in vivo* clearing applications. Second, the compatibility of clearing agents with living tissues depends not only on osmolality but also on their chemical composition. For instance, tartrazine is a trisodium salt; while its optical properties arise primarily from the aromatic anion, the associated three Na^+^ ions may perturb the physiology of electrogenic cells such as neurons and cardiomyocytes, whose resting membrane potentials are highly sensitive to extracellular sodium concentrations. This consideration may limit the applicability of tartrazine trisodium salt in neural tissues, although alternative counterions (e.g., a bio-orthogonal trivalent cation) could mitigate such effects and also reduce osmotic burden while maintaining clearing effect. Finally, the optimal RI for effective *in vivo* clearing is likely tissue-dependent too, reflecting differences in membrane permeability of the clearing agent and the spatial distribution of dominant optical scatterers across cell and tissue types.

This study still has several outstanding questions. One key issue is that the hyperosmolality of tartrazine solutions does not necessarily translate to an equivalent degree of hypertonicity, as tartrazine may cross the cell membrane and therefore may not be fully osmotically active. Although the anionic nature of tartrazine might suggest limited membrane permeability, our unpublished observations indicate the presence of tartrazine in the cytoplasm of cultured cells following brief (∼30 s) exposure, suggesting potential membrane permeability. This finding may explain why we did not observe an optimal RI for live-cell clearing at 1.36∼1.37, but instead observed increasing transparency up to 1.41 (**Fig. 2**). This warrants further investigation to determine the membrane permeability of tartrazine and other absorbing dye molecules. Although still a hypothesis, this interpretation is consistent with the recent BSA-based clearing study, which reported that tartrazine achieved substantial (∼10×) optical clearing of suspended live HeLa cells relative to PBS even at an RI of 1.43 (Ref. 39, Extended Data Fig. 3k), whereas the membrane-impermeable clearing agent iodixanol at the same RI exhibited no measurable clearing effect (Ref. 39, Fig. 1b). Another limitation is that, although the tartrazine/gelatin clearing solution largely preserves cell viability after up to 45 min of continuous exposure, this represents only a temporary solution. An ideal *in vivo* clearing agent should maintain cell viability and function under continuous, long-term exposure. We propose that the Kramers-Kronig relations provide a framework for addressing this challenge by guiding the identification of much more strongly absorbing molecules capable of achieving effective RI modulation at sub-millimolar or even micromolar concentrations, thereby minimizing perturbation to the physiological environment.

## Methods

### Chemicals and materials

Tartrazine (T0388) and gelatin (G6144) were purchased from Sigma-Aldrich. Ultrapure water was obtained using a Millipore Milli-Q Integral 10 water purification system. The silica sphere suspension (SISN1000, 1 μm diameter) was procured from nanoComposix. Other materials, including DMEM (10569010), fetal bovine serum (FBS, A3160802), penicillin-streptomycin (15070063), trypsin-EDTA (25300054), poly-D-lysine (PDL, A3890401), and 1× phosphate-buffered saline (PBS, 20012027), were sourced from Gibco. Additionally, 3-(4,5-dimethylthiazol-2-yl)-2,5-diphenyltetrazolium bromide (MTT, M6494), dimethyl sulfoxide (DMSO, D12345), and the LIVE/DEAD Cell Imaging Kit (R37601) were obtained from Invitrogen. Iodixanol (Cat. No: 1893) was purchased from PROGEN Biotech Inc.

### Ellipsometry of tartrazine and gelatin solutions

Ellipsometry was performed to determine the real and imaginary components of the refractive index of solutions containing varying concentrations of tartrazine and gelatin. Measurements were conducted using a Horiba UVISEL ellipsometer. A shallow vinyl specimen mold (Tissue-Tek Cryomold, Sakura Finetek USA) with defined dimensions (25 mm length × 20 mm width × 5 mm height) was used; its edges were removed, and a sandpaper pad was affixed to the bottom surface. Approximately 2.1 mL of each solution was added to the mold to form a flat, reflective air-liquid interface at the top. The complex refractive index spectrum of each solution was determined using a single-interface, semi-infinite reflection model, with reflected light collected at an incidence angle of 69.85°. An integration time of 0.2 s per wavelength was applied for all measurements.

### UV-Vis transmission and absorption spectroscopy

UV-Vis transmission and absorption spectroscopy were conducted using a Thermo Fisher Scientific Evolution 350 UV-Vis spectrophotometer. An acquisition time of 0.2 s was applied at each wavelength, and all measurements were performed over a wavelength range of 190 to 1100 nm. For transmission and absorbance measurements of the clearing agents, 1-mm and 10-µm pathlength quartz cuvettes were used, respectively, so as to avoid saturation of the measurements. Prior to use, the cuvettes were cleaned by rinsing with aqua regia followed by water, and then dried thoroughly. Reference spectra were collected using water alone to account for optical losses due to intrinsic water absorption and reflections at the interfaces.

### Cell culture

Cells were cultured in a standard humidified incubator at 37 °C with 5% CO_2_. Human embryonic kidney (HEK) cells were routinely maintained in DMEM supplemented with 10% FBS and 1% penicillin-streptomycin. The HEK cell line was acquired from the American Type Culture Collection (ATCC Number: CRL-1573). Culture dishes were pre-coated with poly-D-lysine for 2 h at 37 °C, followed by a PBS wash prior to cell seeding. After seeding onto confocal dishes or culture plates, cells were allowed to proliferate for 24 h before being used in experiments.

### Preparation of tartrazine/gelatin solutions

Tartrazine powder was dissolved in sterile Milli-Q water with varying concentrations of gelatin to prepare several formulations: 0.18 M tartrazine with 5.8% w/w gelatin, 0.33 M tartrazine with 5.8% gelatin, 0.47 M tartrazine with 5.8% gelatin, 0.47 M tartrazine alone, 0.47 M tartrazine with 1.5% gelatin, and 0.47 M tartrazine with 3% gelatin (**Table S1**). The solutions were then heated in an oven (Heratherm OMH60) at 80 °C to facilitate complete dissolution. Afterward, the solutions were allowed to cool to room temperature prior to subsequent testing. Before conducting cellular experiments, the pH of each solution was adjusted to 7.2∼7.4 using a pH meter (SevenDirect SD20).

### Cell imaging experiments

Microscopic imaging of cultured cells was performed using a commercial Olympus microscope equipped with an LED light source and a DIC module. Cultured HEK cells in a confocal dish were positioned under the objective, and the culture medium was removed prior to the addition of a solution containing specified concentrations of tartrazine and gelatin. During imaging, the light intensity was adjusted to compensate for tartrazine’s absorption at blue wavelengths. For reversal, the tartrazine/gelatin solution was carefully removed to maintain a consistent field of view under the objective. Subsequently, 5 mL of 1× PBS was added to the dish as a washout step. Cell conditions were monitored using a camera at a frame rate of 1 Hz.

### Cytotoxicity assessment

For cytotoxicity assessment, HEK cells were seeded in confocal dishes pretreated with poly-D-lysine and incubated for 24 h. After exposure to tartrazine/gelatin solutions for 15, 30, or 45 min, the solutions were aspirated, and the cells were gently washed with PBS. The negative control group did not undergo any exposure to tartrazine/gelatin solutions. As a positive control for toxicity, 20% DMSO was applied under the same conditions. An iodixanol solution with a matched RI of 1.41 was applied under identical conditions for comparison, as it has been identified as an alternative tissue-clearing agent with demonstrated safety in cultured cells.^48^ Cells were then stained using the LIVE/DEAD Cell Imaging Kit and imaged with a fluorescence microscope (Leica DMi8). Cell viability was quantified by calculating the ratio of live cells to total cells, with analysis performed using ImageJ software.

In addition, cell metabolic activity was evaluated using the MTT assay. HEK cells were cultured in 96-well plates and subjected to various treatment conditions. Following incubation with tartrazine/gelatin solutions for different time periods, the solutions were aspirated, and the cells were incubated with 0.5 mg/mL MTT in culture medium for 4 h at 37 °C.^60^ After incubation, the resulting formazan crystals were solubilized in DMSO and agitated for 15 min. The absorbance of the solubilized formazan was measured at 570 nm using a microplate reader (BioTek Synergy H1).

### Flow cytometry analysis

HEK cells cultured in 35 mm dishes were exposed to 0.47 M tartrazine and 5.8% gelatin for 30 minutes. After treatment, cells were rinsed with 1× PBS to remove the clearing solution and subsequently cultured for an additional 48 hours to evaluate the delayed effects of the clearing solution on cell viability. Both control (without treatment) and treated cells were dissociated with 0.05% trypsin-EDTA and neutralized with 1× PBS containing 2% FBS. Cells were harvested by centrifugation, stained with Calcein-AM/BI following the manufacturer’s protocol, and filtered through a 45 μm cell strainer into a 96-well round-bottom plate for immediate analysis on a Beckman Coulter CytoFlex S flow cytometer. Flow cytometry data were processed and analyzed using FlowJo 10 software.

### Image processing

The captured images were processed using ImageJ software. Image contrast was calculated using the following formula:

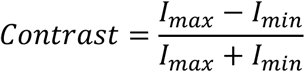

where *I*_*max*_ and *I*_*min*_ represent the maximum and minimum grayscale intensity values, respectively, obtained from each corresponding microscopic image.

To determine cellular size changes, cell areas in DIC images were outlined and quantified using ImageJ. Because HEK cells in culture often form densely packed clusters, it is challenging to accurately delineate individual cell boundaries within these clusters. Instead, the overall outline of each entire cluster was traced to obtain the total area, which was then divided by the number of constituent cells in the cluster to calculate the average cell area. The percentage of shrinkage was determined by dividing the average cell area after treatment by that before treatment.

### Statistical analysis

All statistical analyses and graph preparations were performed using GraphPad Prism software. Two-tailed unpaired *t*-tests and one-way ANOVA were used to assess statistical significance. A *P* value of less than 0.05 was considered statistically significant. The specific statistical tests used for each figure panel are detailed in the corresponding figure legends. Data are presented as mean ± standard deviation (S.D.), as specified.

## Acknowledgments

G.H. acknowledges three awards from the NIH (5R00AG056636-04, 1R34NS127103-01, and R01NS126076), an NSF CAREER award (2045120), an NSF EAGER award (2217582), a Rita Allen Foundation Scholars Award, a Beckman Technology Development Grant, a grant from the Focused Ultrasound Foundation, a gift from the Spinal Muscular Atrophy (SMA) Foundation, two gifts from the Pinetops Foundation, two seed grants from the Wu Tsai Neurosciences Institute, two seed grants from the Bio-X Initiative of Stanford University, a seed grant from a Wu Tsai Synthetic Neuroscience Grant, and a teacher-scholar award from the Camille and Henry Dreyfus Foundation. M.L.B. acknowledges a grant from the Air Force Office of Scientific Research (FA9550-21-1-0312). S.C. acknowledges support from the Stanford Bio-X Bowes Fellowship. H.C. acknowledges support from the Stanford Interdisciplinary Graduate Fellowship.

## Author Contributions

X.H. and G.H. contributed to the conceptualization; X.H. and G.H. contributed to the methodology; X.H., S.C., and Z.L. contributed to the investigation; X.H., H.C., S.Z., L.Z., A.B., and V.C. contributed to the data analysis; X.H., M.L.B., and G.H. contributed to the manuscript preparation; G.H. and M.L.B. contributed to the funding acquisition.

