## Supplementary Information for "Tartrazine clears live cells while preserving viability at high refractive indices and osmolality"

### Table of Contents

Figures S1–5……………………………………………………………………………….2

Tables S1…………………………………………………………………………………..7

Movie captions S1–2………………………………………………………………………8

**Supplementary figures**

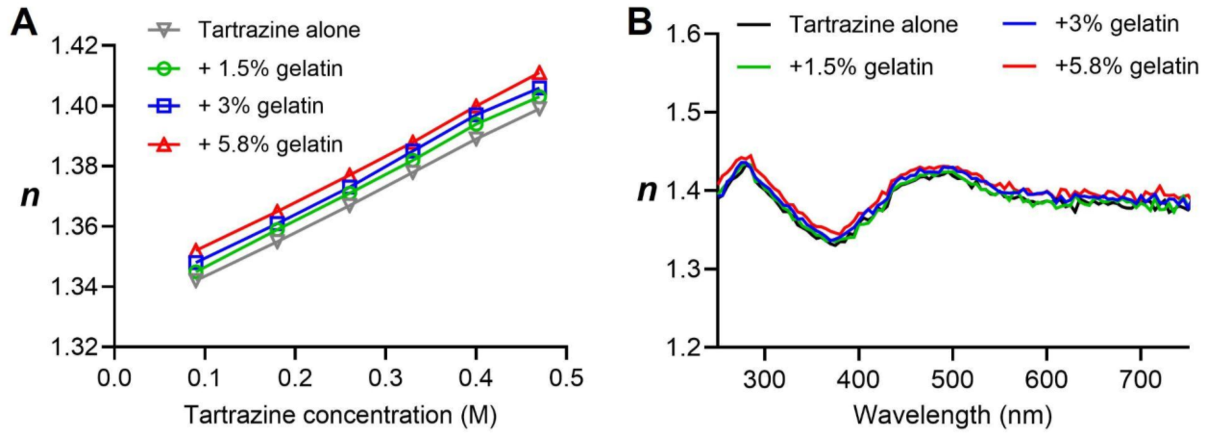


**Figure S1. RI dependence on tartrazine and gelatin concentrations.** (**A**) Dependence of the real refractive index (*n*) measured by an Abbe refractometer on the concentration of tartrazine with different gelatin concentrations. (**B**) *n* of 0.47 M tartrazine solutions containing different concentrations of gelatin measured as a function of wavelength by ellipsometry.

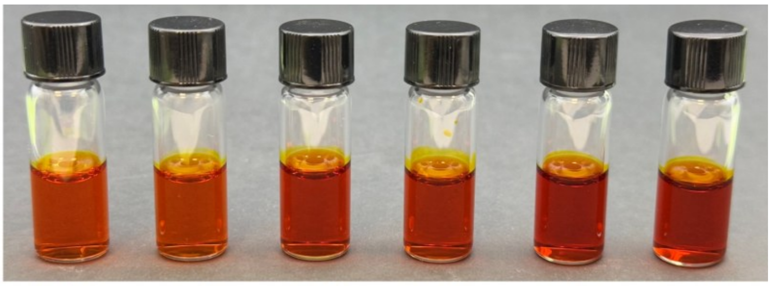

| | | | | | | |
|---|---|---|---|---|---|---|
| Tartrazine (M) | 0.18 | 0.18 | 0.33 | 0.33 | 0.47 | 0.47 |
| Gelatin (% w/w) | 0 | 5.8 | 0 | 5.8 | 0 | 5.8 |

**Figure S2. Photographs of solutions containing varying concentrations of tartrazine and gelatin.**

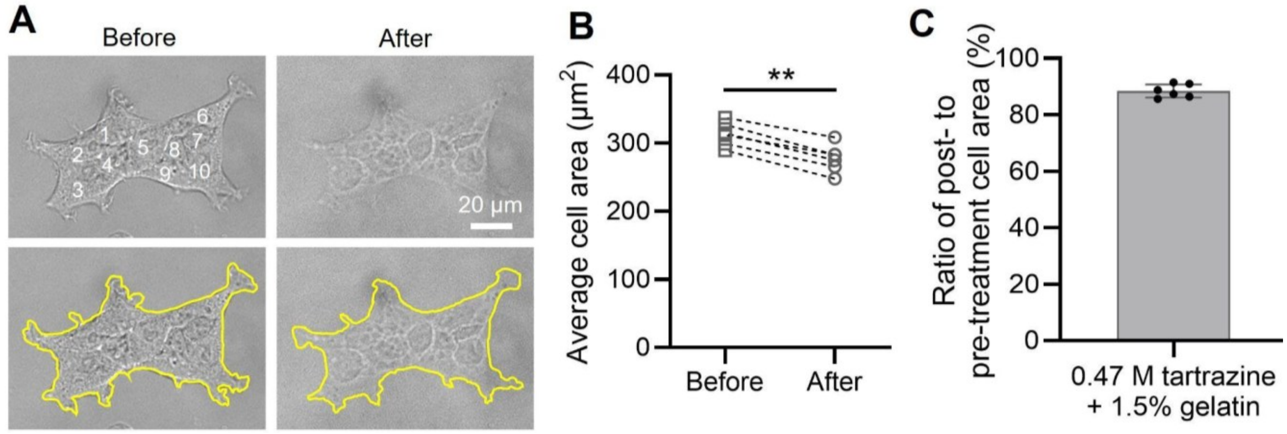


**Figure S3. Cell morphology and size changes in response to the tartrazine solution containing 1.5% gelatin.** (**A**) HEK cells immersed in a solution containing 0.47 M tartrazine and 1.5% w/w gelatin. (**B**) Quantification of the average cell area following tartrazine/gelatin treatment in **A**. (**C**) Quantification of the percent cell area change following tartrazine/gelatin treatment in **A**. $^{**}p < 0.01$, as determined by unpaired two-tailed t-tests. $n = 6$ biological replicates for panels **B** and **C**. Data in panels **C** is presented as mean ± SD.

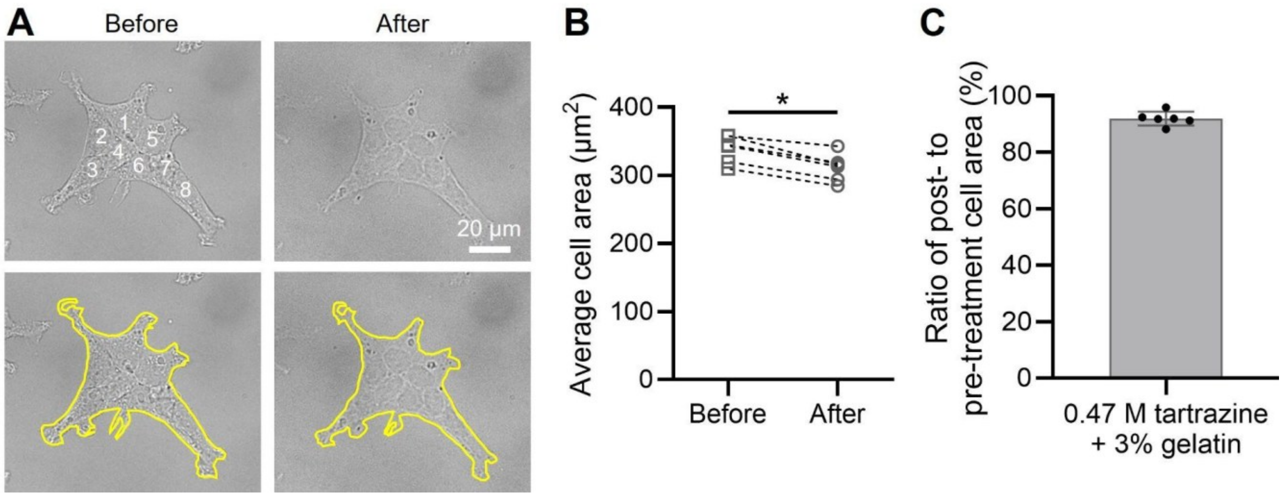


**Figure S4. Cell morphology and size changes in response to the tartrazine solution containing 3% gelatin.** (**A**) HEK cells immersed in a solution containing 0.47 M tartrazine and 3% w/w gelatin. (**B**) Quantification of the average cell area following tartrazine/gelatin treatment in **A**. (**C**) Quantification of the percent cell area change following tartrazine/gelatin treatment in **A**. *$p < 0.05$, as determined by unpaired two-tailed t-tests. $n = 6$ biological replicates for panels **B** and **C**. Data in panels **C** is presented as mean ± SD.

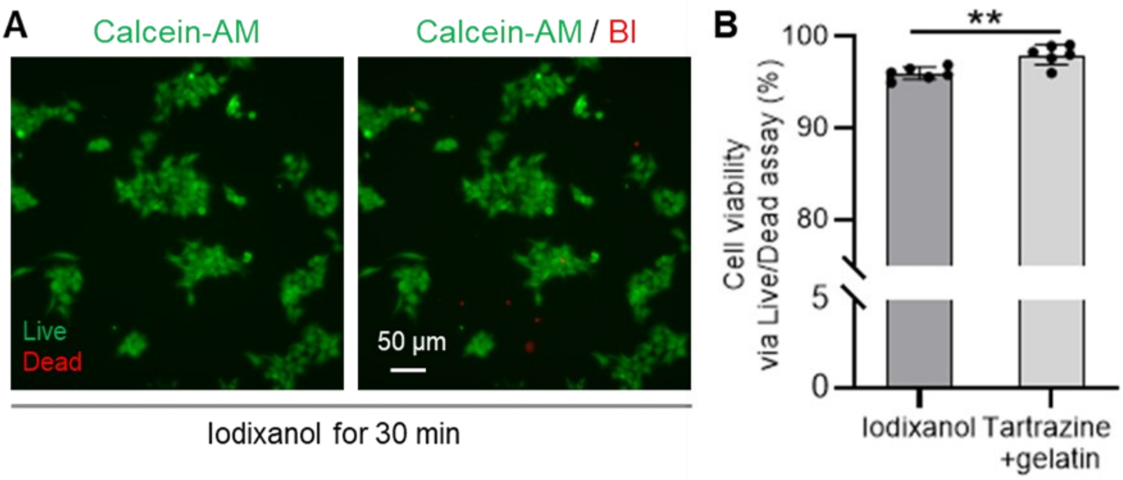


**Figure S5. In vitro cell viability assessment of another clearing agent, iodixanol.** (**A**) Representative images of live/dead staining in HEK cells after 30 minutes of incubation with iodixanol (RI 1.41). Live cells are shown in green, and dead cells in red. (**B**) Quantification of cell viability following incubation, based on live/dead staining. Data from the 30-minute incubation with 0.47 M tartrazine and 5.8% gelatin is also presented in **Fig. 4B**. $^{**}p < 0.01$, as determined by unpaired two-tailed t-tests. Data are presented as mean ± SD in panel **B** for $n = 6$ biological replicates.

**Table S1. Chemical compositions of different tartrazine/gelatin solutions.**

| | Tartrazine (mg) | Water (mg) | Gelatin (mg) |
|---|---|---|---|
| 0.18 M tartrazine + 5.8% gelatin | 100 | 1000 | 67.7 |
| 0.33 M tartrazine + 5.8% gelatin | 200 | 1000 | 73.8 |
| 0.47 M tartrazine + 5.8% gelatin | 300 | 1000 | 80 |
| 0.47 M tartrazine | 300 | 1000 | 0 |
| 0.47 M tartrazine + 1.5% gelatin | 300 | 1000 | 19.8 |
| 0.47 M tartrazine + 3% gelatin | 300 | 1000 | 40.2 |

**Movie captions**

**Movie S1.** HEK cells lose membrane contrast upon addition of 0.47 M tartrazine and 5.8% gelatin, with no significant cell shrinkage observed. The video is shown at 5× real-time speed.

**Movie S2.** HEK cells lose membrane contrast upon addition of 0.47 M tartrazine, resulting in noticeable cell shrinkage. The video is displayed at 5× real-time speed.
